# Squalene as a Candidate Signalling Molecule: Computational Evidence for a Predicted KEAP1 Interaction and Implications for Stress-Induced Altruistic Stem-Cell Niche Protection

**DOI:** 10.64898/2026.11.03.763951

**Authors:** Partheevjyoti Dutta, Upasha Sarmah, Rupam Das, Partha Jyoti Saikia, Lekhika Pathak, Debduti Datta, Seema Bhuyan, Bikul Das

## Abstract

Stem and progenitor cells exposed to cytotoxic stress must preserve both their own viability and the functional integrity of their surrounding niche. Previous investigations from our laboratory have described stress-adapted altruistic stem-cell phenotypes and demonstrated the cytoprotective potential of squalene in normal haematopoietic and mesenchymal progenitor cells through preservation of glutathione homeostasis. More recent studies have associated hypoxia-driven changes in squalene metabolism with intracellular squalene accumulation and protection against lipid peroxidation, while independent experimental evidence has implicated NRF2 in squalene-mediated cytoprotection. However, the molecular basis through which squalene may engage the NRF2 regulatory pathway remains incompletely understood. Here, we investigated the structural feasibility of a potential interaction between squalene and the KEAP1 Kelch domain using molecular docking across three human KEAP1 crystal structures (PDB IDs: 6TYM, 6SP4 and 5CGJ). The docking protocol was evaluated through native-ligand redocking, yielding RMSD values of 0.826, 0.277 and 0.303 Å, respectively. Squalene was consistently accommodated within the canonical hydrophobic cavity, with predicted docking scores of −7.905, −7.513 and −6.756 kcal/mol. The predicted interaction environment was predominantly hydrophobic and involved residues including Arg415, Arg380 and Asn414, without conventional hydrogen-bond formation. These findings support the structural feasibility of squalene accommodation within the KEAP1 Kelch domain and provide a testable hypothesis connecting stress-responsive squalene metabolism with potential KEAP1–NRF2 regulation. However, docking alone does not establish direct binding, physiological accessibility, KEAP1 inhibition or functional NRF2 activation. Further biophysical, molecular-dynamics and cellular investigations are required to establish the biological significance of the predicted interaction.

## 1. Introduction

Stem and progenitor cells reside within specialised cellular niches that regulate their survival, self-renewal, differentiation and regenerative capacity. An important component of niche maintenance is the regulation of cellular redox homeostasis, including the balance between reduced and oxidised glutathione (GSH). Although antioxidant responses are often considered cell-autonomous, the redox state and functional integrity of a stem-cell compartment can also be influenced by interactions between neighbouring cells (Bigarella et al., 2014; Ito & Suda, 2014). This raises an important question in stem-cell biology: can a subset of stressed cells actively contribute to the protection of neighbouring cells and thereby preserve the surrounding niche?

An early line of investigation from our laboratory examined the protective effects of squalene on normal haematopoietic and mesenchymal bone-marrow progenitor cells. Studies reported that squalene could protect these cells against platinum-associated oxidative toxicity while preserving glutathione homeostasis (Das et al., 2003; Das et al., 2008). These observations provided an initial biological basis for considering squalene as a potential contributor to cellular protection beyond its conventional role as an intermediate in cholesterol biosynthesis.

A subsequent conceptual development was the identification of the altruistic stem-cell phenotype. Das et al. (2012) described a stress-responsive cellular state in which a subset of human embryonic stem cells undergoes reversible changes in p53/MDM2 regulation, accompanied by increased HIF-2α activity and glutathione production. These cells release glutathione into their surrounding environment and thereby contribute to the protection of neighbouring cells from stress-associated differentiation or cell death. The altruistic phenotype was proposed as a transient response, with the protective state eventually resolving through restoration of p53 activity and subsequent apoptosis or differentiation. This work introduced the concept that stem-cell protection may involve an active, temporary contribution from a specialised cellular subpopulation rather than relying exclusively on the intrinsic antioxidant capacity of individual cells.

The concept was subsequently extended towards mesenchymal stem cells (MSCs) and the protection of neighbouring cells within hypoxic and cytotoxic environments. In 2017, Pal et al. reported the investigation of BCG-infected mesenchymal stem cells as a strategy for targeting oral cancer stem cells within the hypoxic niche (AACR abstract 3903). This work provided an early link between MSC-mediated responses, pathogen exposure and the regulation of cancer stem-cell populations in a hypoxic microenvironment. It also broadened the scope of niche defence from normal stem-cell compartments towards the tumour microenvironment.

In 2022, Bhuyan et al. further developed this concept through the description of pathogen-induced stem-cell niche defence in hypoxia-induced tumour stemness defence (TSD) cells. Their study demonstrated that BCG infection of TSD-associated cells was linked to HMGB1/p53-related signalling and TLR2/TLR4-dependent bystander apoptosis in neighbouring cancer stem cells. These findings provided a mechanistic framework in which stress-adapted tumour cells could participate in pathogen-associated responses that influence the survival of neighbouring cancer stem cells (Bhuyan et al., 2022). Thus, the concept of niche defence expanded from antioxidant-mediated protection in normal stem-cell systems to pathogen-associated modulation of the tumour stem-cell compartment.

The investigation was further extended towards the relationship between the oral microbiome and tumour stemness defence. Saikia et al. (2023) reported that *Fusobacterium nucleatum* associated with oral cancer subjects could contribute to the reprogramming of oral cancer stem cells towards a TSD-associated phenotype. This work suggested a possible relationship between microbial exposure, tumour-cell plasticity and niche-defence-associated signalling. It provides a further research direction connecting microbial interactions with the regulation of tumour stem-cell states.

In 2025, Pathak et al. described hypoxia-driven mobilisation of altruistic cancer stem cells in platinum-treated head and neck cancer. This work extended the investigation of stress-adapted cellular phenotypes to a clinically relevant treatment-associated setting, further developing the relationship between hypoxia, cytotoxic stress and protective cellular responses in the tumour microenvironment (Pathak et al., 2025). In 2026, Sandhya et al. investigated the stemness, immunomodulatory and biodistribution characteristics of clinical-grade MSCs in relation to niche-defence-associated cytoprotection against platinum-mediated toxicity and coronavirus infection. In the same year, Talukdar et al. further discussed adaptive altruism and dormant cancer stem-cell niche defence under hypoxic conditions (Sandhya et al., 2026; Talukdar et al., 2026). Collectively, these studies represent a progression from the initial observation of stress-associated progenitor-cell protection towards the investigation of specialised cellular phenotypes, intercellular protection, pathogen responses and translational applications of stem-cell niche defence.

Despite these advances, the metabolic mechanisms that support stress-adapted cytoprotection and maintain antioxidant capacity within the surrounding niche remain incompletely understood. In particular, the relationship between the glutathione-associated protective response and the availability of other stress-responsive metabolites warrants further investigation. In this context, squalene is of interest because of its earlier association with the protection of normal haematopoietic and mesenchymal progenitors against platinum-induced oxidative toxicity (Das et al., 2003; Das et al., 2008). In parallel, laboratory observations in patient-derived TSD-positive cellular systems have identified extracellular squalene under hypoxic conditions. These observations provide a rationale for investigating squalene in the context of niche defence; however, the reported glutathione-associated secretory response and the parallel observation of extracellular squalene should be treated as distinct experimental findings. The latter should not be attributed to previous studies unless directly demonstrated in those studies.

Squalene is a 30-carbon, linear, uncharged triterpene synthesised through the mevalonate pathway and serves as an intermediate in sterol biosynthesis (Smith, 2000; Spanova & Daum, 2011). Its highly hydrophobic and conformationally flexible structure favours partitioning into lipid-rich environments. Importantly, cellular squalene is not necessarily directed entirely towards cholesterol synthesis. Studies in human adipocytes have demonstrated distinct metabolic pools of squalene, including a rapidly turning-over pool and a larger storage pool, suggesting that its intracellular availability is subject to metabolic compartmentalisation (Tilvis et al., 1982). This raises the possibility that squalene availability may be influenced by cellular metabolic state and may have biological consequences beyond its role as a sterol precursor.

Recent studies have provided further support for a relationship between squalene metabolism and cellular stress adaptation. Hypoxia-associated changes in squalene monooxygenase (SQLE), the oxygen-dependent enzyme responsible for converting squalene into 2,3-oxidosqualene, can promote intracellular squalene accumulation (Garcia-Bermudez et al., 2019; Coates et al., 2023). Squalene accumulation associated with reduced SQLE activity has also been linked to changes in membrane lipid composition, reduced lipid peroxidation and decreased susceptibility to ferroptotic injury (Garcia-Bermudez et al., 2019). These findings suggest that hypoxia-associated changes in squalene metabolism may contribute to cellular protection through membrane-associated mechanisms. Whether squalene also participates in protein-mediated antioxidant signalling remains an important question.

Independent experimental evidence has further connected squalene with NRF2-dependent cytoprotection. Jiang et al. (2026) reported that squalene alleviated experimental colitis through restoration of NRF2-associated redox balance. The protective effects were attenuated by genetic deletion or pharmacological inhibition of NRF2, while cellular thermal shift assays and complementary computational analyses suggested potential molecular engagement. These findings support a role for NRF2 in squalene-associated cytoprotection in that experimental model. However, the precise molecular mechanism through which squalene engages the NRF2 regulatory system remains to be fully established.

The Kelch-like ECH-associated protein 1 (KEAP1)–NRF2 pathway is a central regulator of cellular antioxidant and cytoprotective responses. Under basal conditions, KEAP1 acts as a substrate adaptor for Cullin-3-dependent ubiquitination of NRF2, promoting its proteasomal degradation (Itoh et al., 1999; Kobayashi et al., 2004). Disruption of KEAP1-mediated regulation allows NRF2 stabilisation and the transcriptional activation of antioxidant-response genes, including genes involved in glutathione biosynthesis and redox regulation, such as *GCLC*, *GCLM* and *SLC7A11* (Baird & Yamamoto, 2020; Cuadrado et al., 2019). Given the independent evidence for NRF2-dependent squalene-mediated cytoprotection, investigating whether squalene can interact with components of the KEAP1–NRF2 regulatory axis may provide a possible molecular link between squalene metabolism and antioxidant responses.

The KEAP1 Kelch domain contains the protein-interaction surface responsible for recognising the ETGE and DLG motifs of NRF2. Squalene, in contrast, is an uncharged and highly hydrophobic molecule with an extended, flexible hydrocarbon backbone. These physicochemical properties provide a rationale for examining whether squalene can be computationally accommodated within the hydrophobic ligand-binding cavity of the KEAP1 Kelch domain. We therefore hypothesised that squalene may exhibit structural compatibility with this domain, providing a testable possibility for interaction with the KEAP1–NRF2 regulatory system. This hypothesis does not assume that NRF2 drives squalene biosynthesis. Rather, hypoxia-associated changes in SQLE activity and mevalonate-pathway metabolism are considered potential determinants of squalene availability, while KEAP1–NRF2 is considered a candidate downstream regulatory pathway.

In the present study, we investigated the structural feasibility of a potential squalene–KEAP1 interaction using molecular docking across three human KEAP1 Kelch-domain crystal structures (PDB IDs: 6TYM, 6SP4 and 5CGJ). Native-ligand redocking was performed to evaluate the ability of the docking protocol to reproduce crystallographic ligand poses, followed by comparative docking and residue-interaction analysis of squalene. The objective was to determine whether squalene could be computationally accommodated within the canonical KEAP1 Kelch-domain cavity, rather than to establish direct binding, KEAP1 inhibition or functional NRF2 activation. The findings provide a structural hypothesis for further investigation of the relationship between stress-responsive squalene metabolism, KEAP1–NRF2-associated antioxidant regulation and stem-cell niche defence.

## 2. Materials and Methods

### 2.1 Study Design

The present study was designed to investigate the structural feasibility of squalene accommodation within the KEAP1 Kelch domain using a structure-based computational approach. Based on previous experimental observations linking squalene with glutathione homeostasis and stem/progenitor-cell protection, together with recent evidence that squalene exerts NRF2-dependent cytoprotective effects, we examined the complementary hypothesis that squalene may also be structurally accommodated within the KEAP1 Kelch domain, the principal protein-interaction domain regulating NRF2 stability.

To investigate this hypothesis, the study was conducted in four sequential stages. First, molecular docking was performed using the Molecular Operating Environment (MOE) to identify potential protein targets and evaluate the interaction of squalene with the KEAP1 Kelch domain (Molecular Operating Environment [MOE], 2019). Second, the docking protocol was independently validated using AutoDock Vina through native ligand redocking to assess the reliability of the docking methodology (Trott & Olson, 2010; Morris et al., 2009). Third, protein–ligand interactions were analysed to identify key amino acid residues contributing to ligand stabilisation and to compare the predicted binding mode of squalene with that of the crystallographic ligand (BIOVIA, Dassault Systèmes, 2021). Finally, the workflow was used to formulate a structural hypothesis for subsequent molecular dynamics analysis and experimental validation of the predicted protein–ligand complex.

This integrative workflow was designed not only to identify KEAP1 as a potential molecular target of squalene but also to provide a mechanistic hypothesis linking proposed squalene-mediated antioxidant activity with regulation of cellular redox homeostasis.

**Figure 1.**
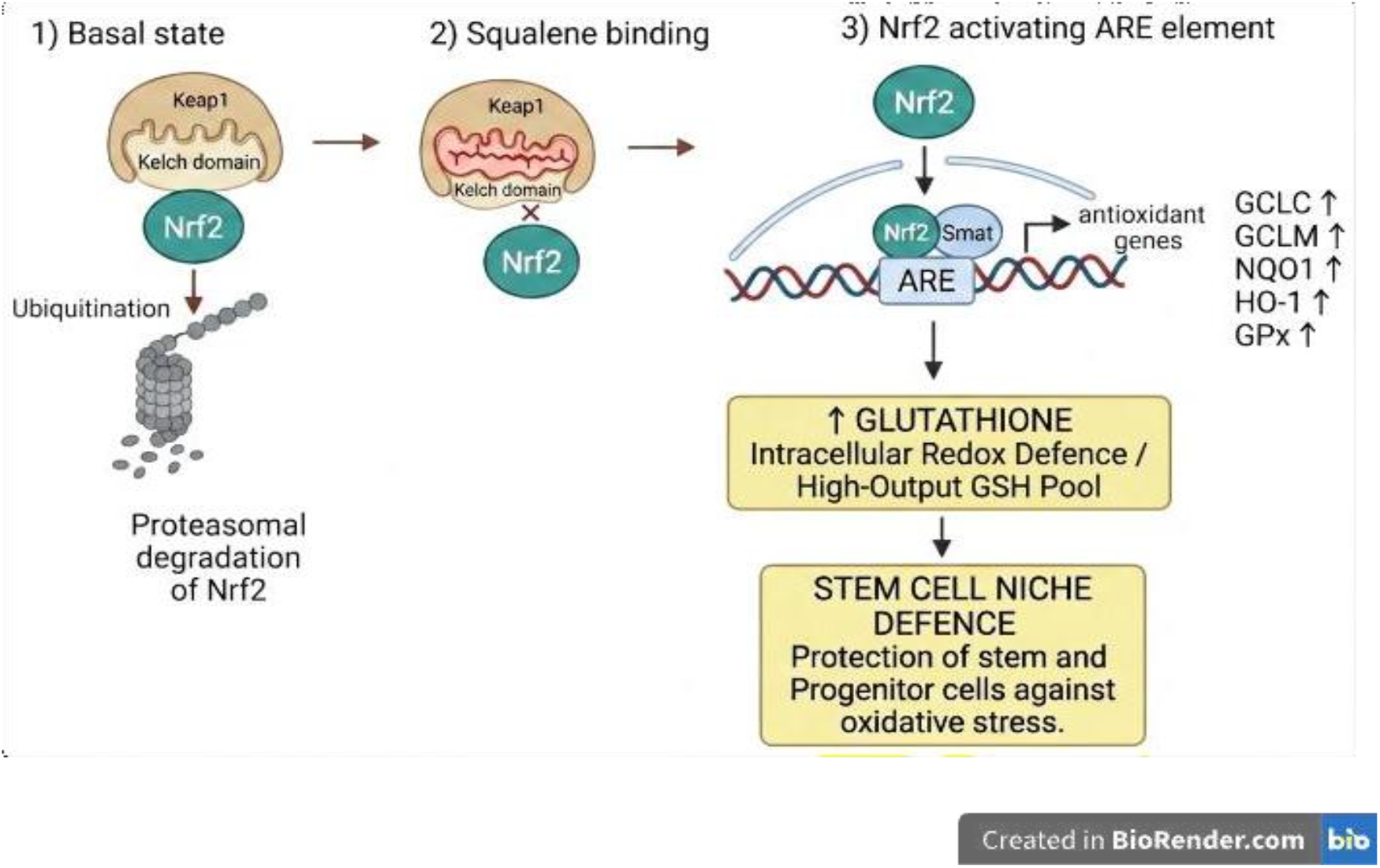
Conceptual framework linking stress-responsive squalene metabolism to stem-cell niche defence. Hypoxia-associated suppression of SQLE can promote squalene accumulation, which may contribute to membrane protection and is proposed here as a candidate signalling metabolite. The predicted KEAP1 interaction provides a testable hypothesis for potential influence on NRF2-dependent antioxidant responses and glutathione-associated cytoprotection; these downstream effects remain to be experimentally established.

**Figure 2.**
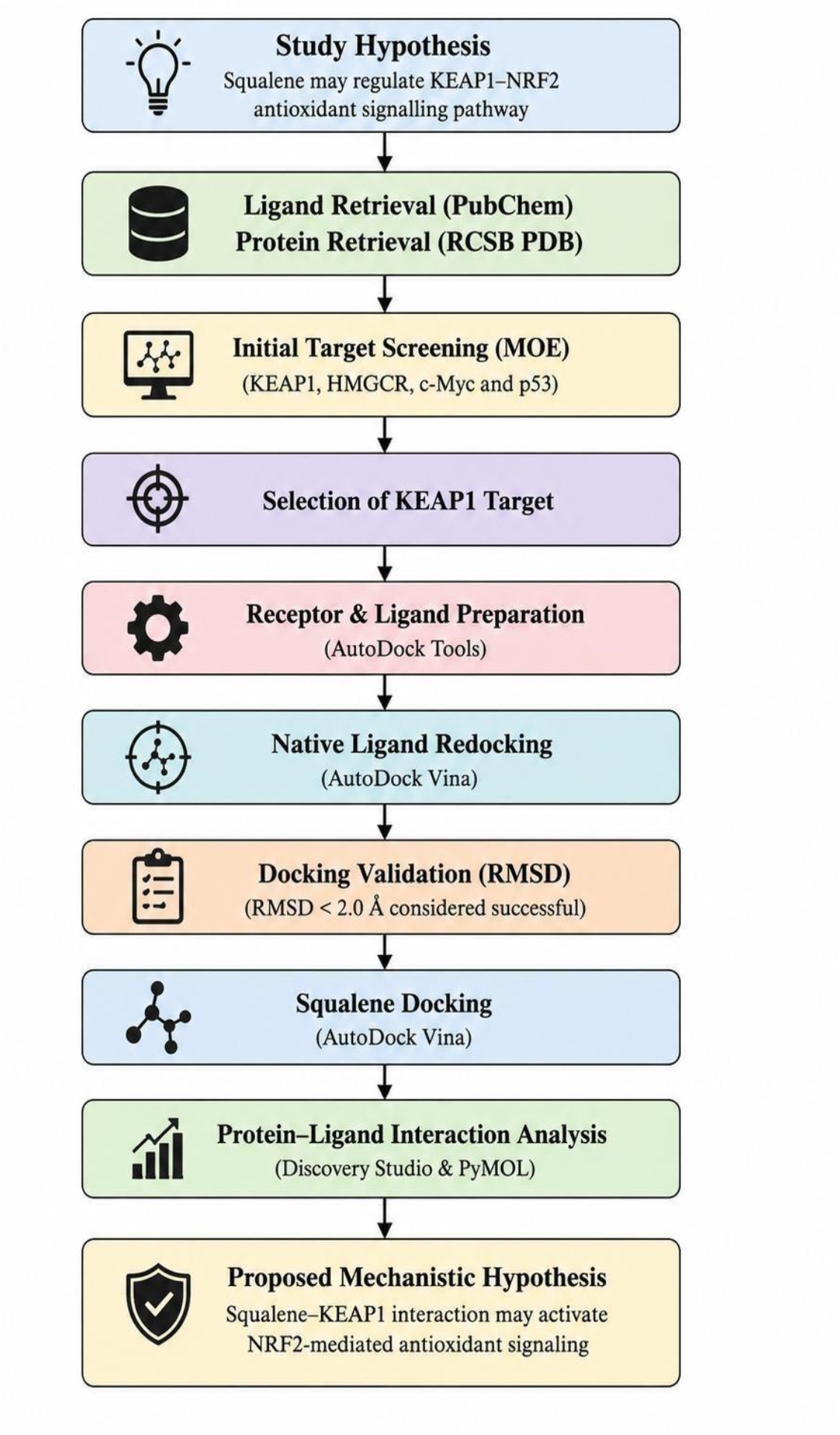
Overview of the computational workflow employed to investigate the interaction between squalene and the KEAP1 Kelch domain. Squalene and protein structures were retrieved from the PubChem and RCSB Protein Data Bank databases, respectively. Initial target screening was performed using Molecular Operating Environment (MOE), followed by selection of KEAP1 for detailed investigation. Receptor and ligand preparation were carried out using AutoDock Tools, and the docking protocol was validated by native ligand redocking using AutoDock Vina. Following successful validation (RMSD < 2.0 Å), squalene was docked into the validated KEAP1 structures. Protein–ligand interactions were subsequently analysed using Discovery Studio Visualizer and PyMOL to propose a mechanistic hypothesis for KEAP1–NRF2-mediated antioxidant signalling.

### 2.2 Selection of Protein Structure

The three-dimensional crystal structures of the human KEAP1 Kelch domain were retrieved from the Protein Data Bank (PDB) (Berman et al., 2000). Multiple crystal structures containing co-crystallised inhibitors were initially evaluated to ensure reproducibility of the docking protocol and to minimise structural bias arising from the use of a single receptor conformation (Ferreira et al., 2015; Pagadala et al., 2017).

Among the available crystal structures, PDB IDs 6TYM, 6SP4, and 5CGJ were selected based on the following criteria:

- human KEAP1 Kelch domain;
- high crystallographic resolution;
- presence of a co-crystallised ligand occupying the canonical NRF2-binding pocket (Jnoff et al., 2014; Jiang et al., 2016);
- suitability for molecular docking validation by native ligand redocking (Trott & Olson, 2010; Morris et al., 2009).

Following protocol validation, the 6TYM crystal structure was selected as the primary receptor for detailed interaction analysis because it demonstrated excellent redocking accuracy and preserved the structural integrity of the canonical KEAP1 ligand-binding cavity.

### 2.3 Protein Preparation

For molecular docking in MOE, the crystal structures of the KEAP1 Kelch domain were prepared by removing crystallographic water molecules and non-essential heteroatoms while retaining the native ligand for binding-site identification. Missing atoms were repaired where necessary, hydrogen atoms were added using the Protonate3D module, and the structures were energy minimised using the AMBER10:EHT force field (Molecular Operating Environment [MOE], 2019; Labute, 2009).

For docking validation and redocking using AutoDock Vina, receptor preparation was performed independently using AutoDock Tools (Morris et al., 2009; Trott & Olson, 2010). The native co-crystallised ligand and crystallographic water molecules were removed, polar hydrogen atoms were added, Gasteiger united-atom charges were assigned, and the receptor structures were converted to PDBQT format (Morris et al., 2009). The docking grid was subsequently centred on the coordinates of the native ligand to enable accurate redocking into the experimentally validated binding pocket (Trott & Olson, 2010)

### 2.4 Ligand Preparation

The three-dimensional structure of squalene (PubChem CID: 638072) was retrieved from the PubChem database in Structure Data File (SDF) format (Kim et al., 2023). For molecular docking using Molecular Operating Environment (MOE), the ligand was imported into MOE, hydrogen atoms were added, and geometry optimization was performed using the AMBER10:EHT force field (Molecular Operating Environment [MOE], 2019). The optimized ligand was subsequently used for docking within the MOE environment.

For docking validation and redocking studies performed using AutoDock Vina, squalene was prepared independently using AutoDock Tools (Morris et al., 2009; Trott & Olson, 2010). The optimized ligand was converted to PDBQT format, Gasteiger partial charges were assigned, non-polar hydrogen atoms were merged, and rotatable bonds were defined according to AutoDock Vina requirements (Morris et al., 2009). The final squalene PDBQT contained 15 torsional degrees of freedom (TORSDOF = 15), and AutoDock Vina was run with an exhaustiveness of 64 to provide broader conformational sampling of the highly flexible ligand. The ligand’s extensive conformational flexibility was retained during docking to allow sampling of alternative conformations within the hydrophobic ligand-binding cavity of KEAP1. Because squalene possesses an extended acyclic polyisoprenoid backbone with substantially greater conformational flexibility than the relatively constrained native ligands used for docking validation, this flexibility was considered explicitly when interpreting the docking results. The squalene-specific AutoDock Vina exhaustiveness value was set to 64 and is reported here to make the conformational sampling protocol reproducible.

**Figure 3.**
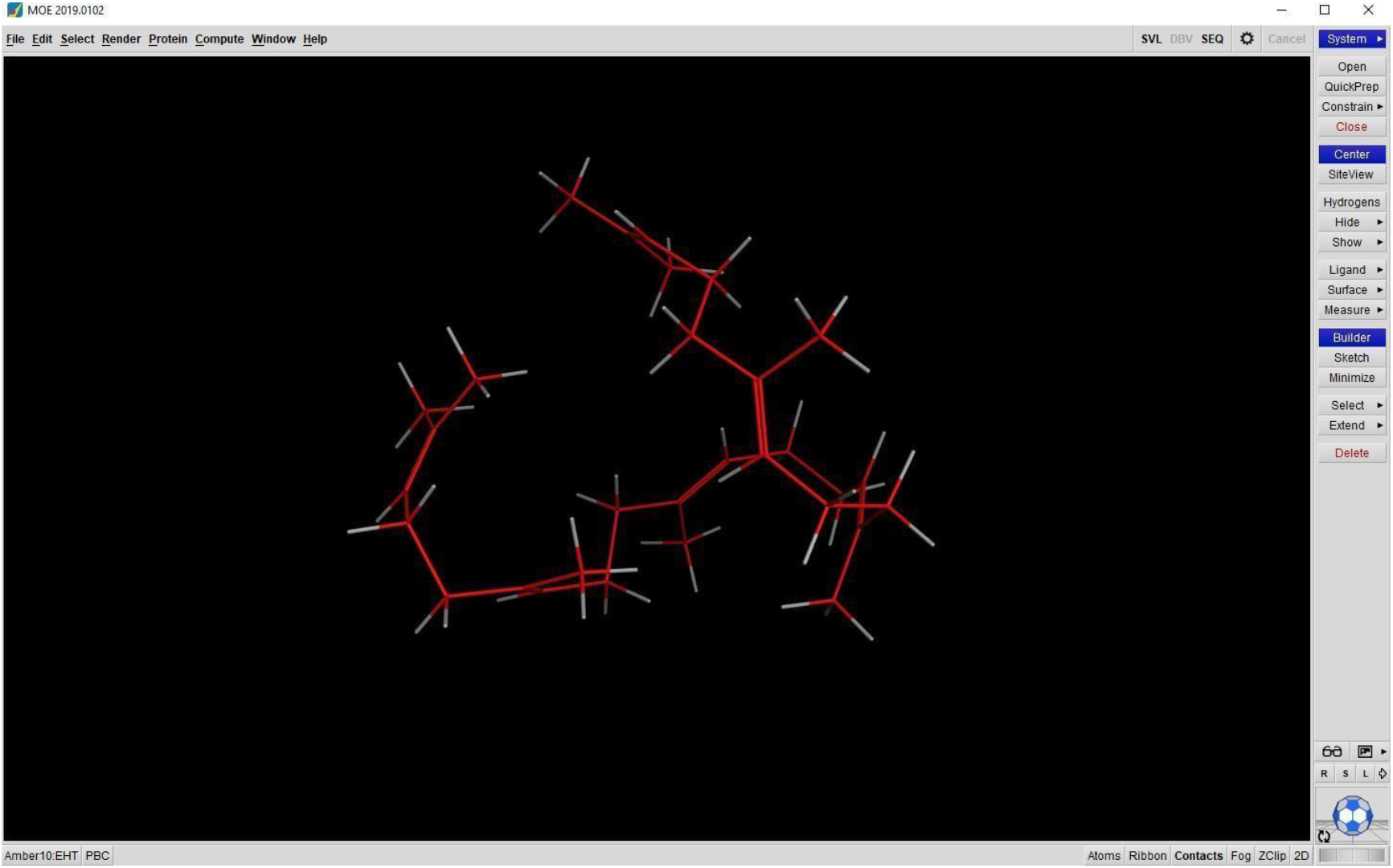
Energy-minimised three-dimensional structure of squalene generated using Molecular Operating Environment (MOE). Hydrogen atoms were added and geometry optimisation was performed using the AMBER10:EHT force field prior to molecular docking. The optimised ligand structure was subsequently used for target screening in MOE and converted to PDBQT format for docking studies using AutoDock Vina.

### 2.5 Molecular Docking

Initial target screening was performed using the Molecular Operating Environment (MOE, Chemical Computing Group, Montreal, Canada) to evaluate selected potential molecular targets of squalene associated with oxidative stress, cellular proliferation, and metabolic regulation (Molecular Operating Environment [MOE], 2019). The selected targets included KEAP1, HMG-CoA reductase, c-Myc, and p53. Protein binding pockets were identified using the **Site** Finder algorithm, and molecular docking was performed using the Triangle Matcher placement method. Initial docking poses were scored using the London dG scoring function, followed by refinement using the Induced Fit protocol and rescoring with the GBVI/WSA dG scoring function (Molecular Operating Environment [MOE], 2019; Labute, 2008). The protein producing the most favourable docking score within this selected set was prioritised for subsequent analysis. This screening was used for target prioritisation within the applied computational workflow and was not intended to provide a quantitative comparison of binding affinities across different proteins or binding sites.

Subsequent molecular docking studies were performed using AutoDock Vina v1.2.7, with receptor and ligand preparation carried out using AutoDock Tools v1.5.7 (Trott & Olson, 2010; Morris et al., 2009). For the initial MOE target-screening stage, the KEAP1 structure represented by PDB ID 1U6D was used for exploratory docking and target prioritisation. This preliminary screening step was distinct from the subsequent AutoDock Vina validation and squalene docking studies, which were performed using the independently selected KEAP1 crystal structures 6TYM, 6SP4, and 5CGJ. Native ligand redocking was first conducted for the selected KEAP1 crystal structures (PDB IDs: 6TYM, 6SP4, and 5CGJ) to validate the docking protocol (Trott & Olson, 2010). Following successful validation, squalene was docked into the canonical ligand-binding pocket of each validated KEAP1 structure using receptor-specific grid definitions centred on the crystallographic ligand coordinates and an exhaustiveness of 64. Docking grids were centred on the coordinates of the co-crystallised ligand to ensure docking within the experimentally validated binding site (Morris et al., 2009). For native-ligand validation, exhaustiveness values of 8 and 32 were used for the 6TYM and 6SP4/5CGJ structures, respectively. For subsequent squalene docking, the exhaustiveness was increased to 64 for all three receptor structures to provide broader conformational sampling of the highly flexible ligand (Trott & Olson, 2010). The docking pose exhibiting the most favourable predicted docking score and a plausible orientation within the canonical KEAP1 ligand-binding pocket was selected for subsequent structural and interaction analyses.

### 2.6 Docking Validation

The reliability of the molecular docking protocol was evaluated by native ligand redocking using AutoDock Vina v1.2.7 (Eberhardt et al., 2021; Trott & Olson, 2010). The co-crystallised ligands from the selected KEAP1 Kelch domain crystal structures (PDB IDs: 6TYM, 6SP4, and 5CGJ) were extracted from their respective Protein Data Bank files and prepared independently using AutoDock Tools v1.5.7 (Berman et al., 2000; Morris et al., 2009). Corresponding receptor structures were prepared by removing the native ligand and crystallographic water molecules, followed by the addition of polar hydrogen atoms, assignment of Gasteiger united-atom charges, and conversion to PDBQT format (Morris et al., 2009).

Redocking was performed by centering the docking grid on the coordinates of the crystallographic ligand to reproduce the experimentally determined binding site. The validated docking protocol was subsequently applied to squalene docking, with the squalene-specific exhaustiveness increased to 64 to account for its greater conformational flexibility (Trott & Olson, 2010). Docking accuracy was assessed by calculating the root mean square deviation (RMSD) between the crystallographic ligand conformation and the corresponding redocked pose using PyMOL (Schrödinger, LLC, 2024). An RMSD value below 2.0 Å was used as a conventional criterion for successful reproduction of the crystallographic ligand pose (Cole et al., 2005), while recognising that RMSD alone does not fully capture interaction-based docking accuracy.

### 2.7 Protein–Ligand Interaction Analysis

The top-ranked docking pose of squalene was analysed using Discovery Studio Visualizer (BIOVIA, Dassault Systèmes, 2021). Three-dimensional visualisation was performed to examine ligand orientation within the KEAP1 binding pocket, whereas two-dimensional interaction diagrams were generated to identify amino acid residues contributing to ligand recognition (BIOVIA, Dassault Systèmes, 2021).

Protein–ligand interactions including hydrophobic contacts, van der Waals interactions, alkyl interactions, π-alkyl interactions, hydrogen bonds, and other non-covalent interactions were identified and compared with those observed for the crystallographic ligand (Ferreira et al., 2015; Guedes et al., 2014). The interaction profile was further evaluated to characterise the predicted interaction environment of squalene within the predefined canonical KEAP1 ligand-binding pocket.

#### Computational resources

Computational docking analyses were performed using an in-house workstation at KaviKrishna Laboratory equipped with an Intel Core i9-12900KF CPU, 32 GB RAM, and an NVIDIA GeForce GTX 1660 Super GPU with 6 GB VRAM. The computational environment was configured using Windows Subsystem for Linux 2 (WSL2) with Ubuntu, and AutoDock Vina v1.2.7 was used for molecular docking. The NVIDIA GPU was available as part of the in-house computational infrastructure; standard AutoDock Vina docking calculations were performed using the Vina computational engine and CPU resources.

### 2.8 Biological Context: Squalene Secretome (Supplementary Table S1)

To provide biological context for the computational docking studies, endogenous squalene and glutathione secretion under cytotoxic stress was examined using a specialized serum-free, hypoxic (1% O_2_) two-phase suspension platform and a two-stage non-contact Boyden chamber assay. The supplementary workflow describes primary human bone marrow mononuclear cell isolation, immunomagnetic reconstitution of CD133^+^/CD271^+^ populations, long term culture –initiating cell (LTC-IC) assessment, cisplatin challenge, and secretome quantification by GC-MS/HPLC. The detailed procedures and reported biological observations are provided in Supplementary Table S1. These data are presented as biological context for the docking hypothesis; they are not a direct experimental test of squalene-KEAP1 binding.

## 3. Results

### 3.1 Identification of KEAP1 as a Potential Molecular Target of Squalene

To explore potential molecular targets associated with the antioxidant activity of squalene, an initial structure-based molecular docking analysis was performed using the Molecular Operating Environment (MOE) (Molecular Operating Environment [MOE], 2019). Squalene was docked against a selected set of proteins implicated in oxidative stress regulation, cellular proliferation, and metabolic signalling, including KEAP1, HMG-CoA reductase, c-Myc, and p53. Among the proteins evaluated under the applied MOE workflow, the KEAP1 Kelch domain produced the most favourable docking score.

The docking scores obtained from the MOE analysis are summarised in Table 1. Squalene produced the most favourable MOE docking score for KEAP1 (S score = −9.74 kcal/mol), followed by HMG-CoA reductase (−8.68 kcal/mol), c-Myc (−8.42 kcal/mol), and p53 (−7.25 **kcal/mol)**. These scores were used to prioritise KEAP1 for subsequent analysis within the scope of the selected protein set and applied docking workflow. Because docking scores obtained for different proteins and binding sites are method-dependent and are not necessarily directly comparable as quantitative measures of binding affinity, the target-screening analysis was considered a prioritisation step rather than definitive evidence of preferential binding. KEAP1 was therefore selected for subsequent docking validation and detailed protein–ligand interaction analysis.

**Table 1.** Preliminary MOE docking scores for squalene against selected protein targets. The KEAP1 target was represented by PDB ID 1U6D. Squalene was evaluated against a selected set of proteins associated with antioxidant defence, lipid metabolism, cell proliferation, and cellular stress responses. The KEAP1 Kelch domain produced the most favourable docking score within the selected protein set under the applied MOE workflow and was therefore prioritised for subsequent docking validation and structural interaction analysis. MOE GBVI/WSA dG scores and AutoDock Vina docking scores are generated by different scoring functions and should not be interpreted as being on a common quantitative scale.

| **Target Protein** | **Docking Score (MOE)** | **Biological Significance** |
|---|---|---|
| KEAP1 Kelch domain<br>*(PDB: 1U6D)* | **−9.74** | Antioxidant signalling |
| HMG-CoA reductase<br>*(PDB ID: 1DQ8)* | −8.68 | Cholesterol biosynthesis |
| c-Myc<br>*(PDB ID: 1NKP)* | −8.42 | Cell proliferation |
| p53<br>*(PDB ID: 1TUP)* | −7.25 | Stress response |

**Figure 4.**
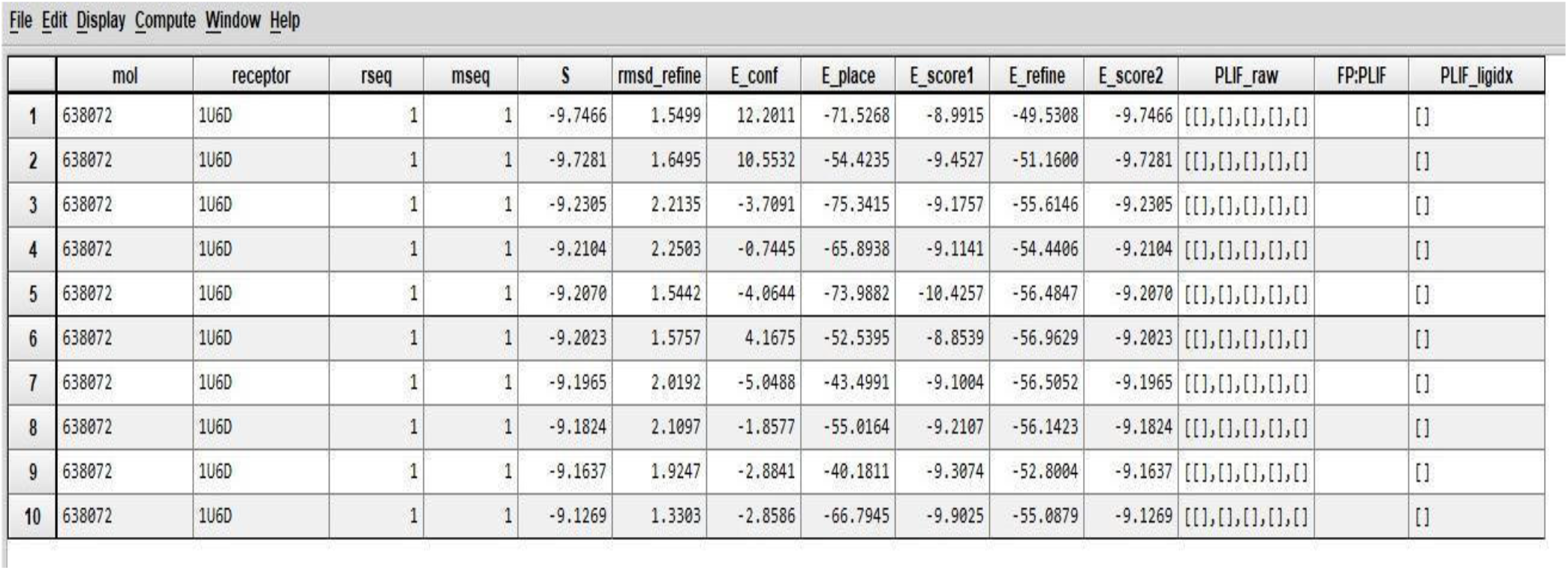
Database Viewer output from Molecular Operating Environment (MOE) showing the ranked docking poses of squalene within the KEAP1 Kelch domain (PDB ID: 1U6D). Docking poses were ranked according to the S score, and the top-ranked pose (Pose 1; S = −9.7466 kcal/mol) was selected for subsequent structural analysis and target prioritisation.

### 3.2 Validation of the Molecular Docking Protocol

Before investigating the interaction between squalene and KEAP1, the docking protocol was validated through native ligand redocking using AutoDock Vina (Trott & Olson, 2010). Three independent human KEAP1 Kelch domain crystal structures containing co-crystallised ligands (PDB IDs: 6TYM, 6SP4, and 5CGJ) were selected for validation (Berman et al., 2000).

The crystallographic ligands were extracted from their respective structures, prepared independently, and redocked into the corresponding ligand-binding cavities using receptor-specific grid definitions centered on the crystallographic ligand coordinates. Docking accuracy was evaluated by calculating the root mean square deviation (RMSD) between the crystallographic ligand conformation and the corresponding redocked pose (Cole et al., 2005; Trott & Olson, 2010).

All three crystal structures exhibited excellent agreement between the experimental and predicted ligand conformations, with RMSD values substantially below the commonly accepted validation threshold of 2.0 Å (Cole et al., 2005) (Table 2). The 6SP4 structure demonstrated the highest accuracy with an RMSD of 0.277 Å, while 6TYM and 5CGJ yielded RMSD values of 0.826 Å and 0.303 Å. These findings support that the docking protocol accurately reproduced the experimentally observed binding mode and was therefore considered suitable for subsequent docking of squalene.

**Table 2:** Native ligand redocking was performed for three experimentally determined KEAP1 Kelch domain crystal structures to validate the docking protocol. Validation was assessed by comparing the crystallographic and redocked ligand conformations using root mean square deviation (RMSD). All structures achieved RMSD values below 2.0 Å, supporting excellent reproduction of the experimental binding modes and the reliability of the docking protocol for subsequent squalene docking studies.

| **PDB ID** | **Native ligand** | **Best redocking score (kcal/mol)** | **RMSD (Å)** | **Status** |
|---|---|---|---|---|
| 6TYM | 08A | **−11.13** | **0.826** | Excellent |
| 6SP4 | LQK | **−11.31** | **0.277** | Excellent |
| 5CGJ | 51M | **−11.04** | **0.303** | Excellent |

**Figure 5.**
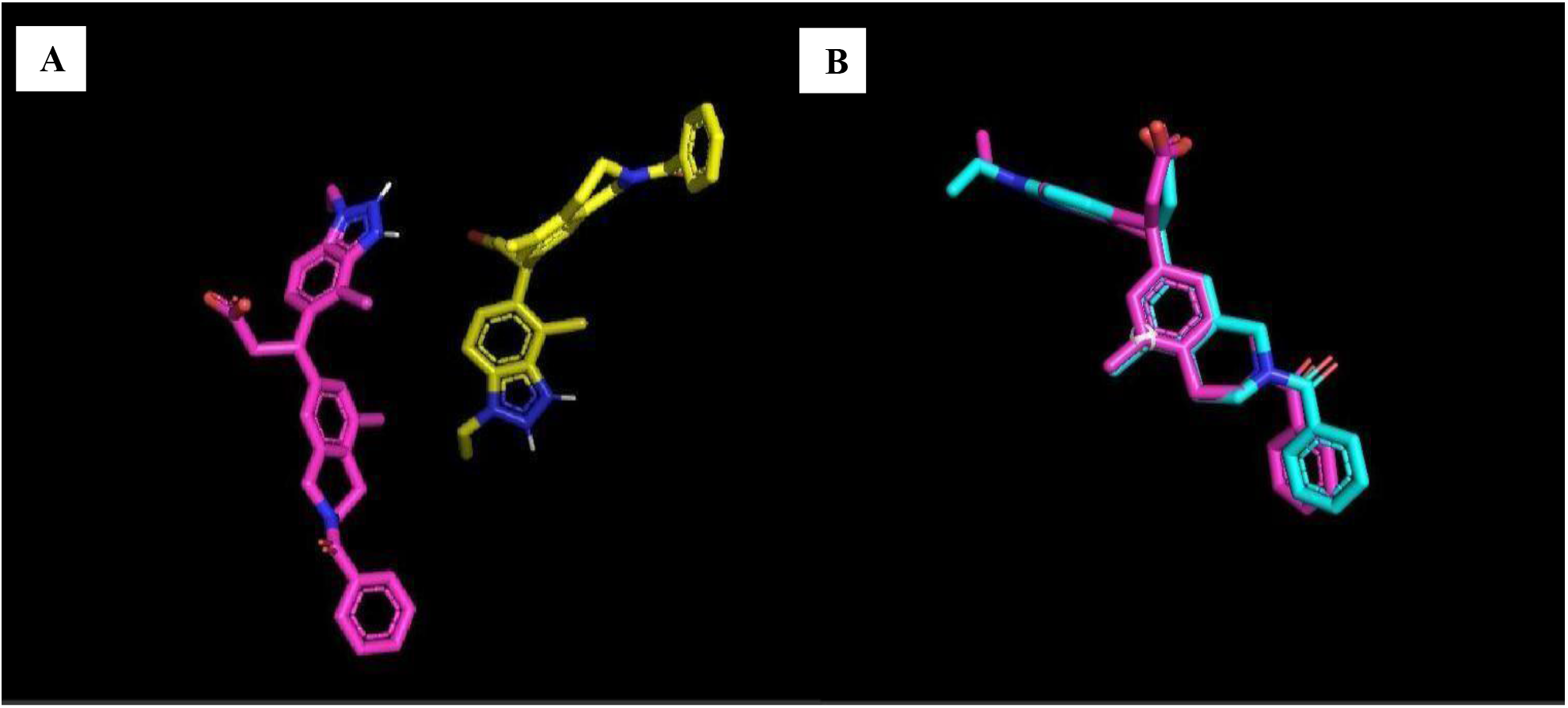
Native ligand (08A) redocking validation of the KEAP1 Kelch domain (PDB ID: 6TYM). **(A)** Experimental binding conformation of the co-crystallised native ligand **08A** (yellow) within the KEAP1 Kelch domain. **(B)** Superposition of the crystallographic ligand **08A** (cyan) and the corresponding redocked pose generated using AutoDock Vina (magenta). The close overlap between the two conformations (RMSD = **0.826 Å**) demonstrates successful reproduction of the experimental binding mode and validates the docking protocol for subsequent squalene docking studies.

### 3.3 Molecular Docking of Squalene with the KEAP1 Kelch Domain

Following successful protocol validation, squalene was docked into the canonical ligand-binding cavity of the KEAP1 Kelch domain. The highest-ranked docking pose positioned squalene within the predefined hydrophobic binding pocket previously occupied by the crystallographic inhibitor.

Despite lacking polar functional groups capable of forming hydrogen bonds or ionic interactions, squalene adopted a predicted conformation compatible with the geometry of the ligand-binding cavity. The predicted binding orientation suggested that ligand stabilisation was primarily mediated through extensive hydrophobic packing and van der Waals interactions distributed along the elongated carbon backbone of the molecule.

Three-dimensional visualisation further demonstrated that squalene was predicted to occupy the canonical NRF2-binding pocket of KEAP1, providing a structurally plausible basis for further investigation of a potential interaction at this regulatory site.

**Table 3:**
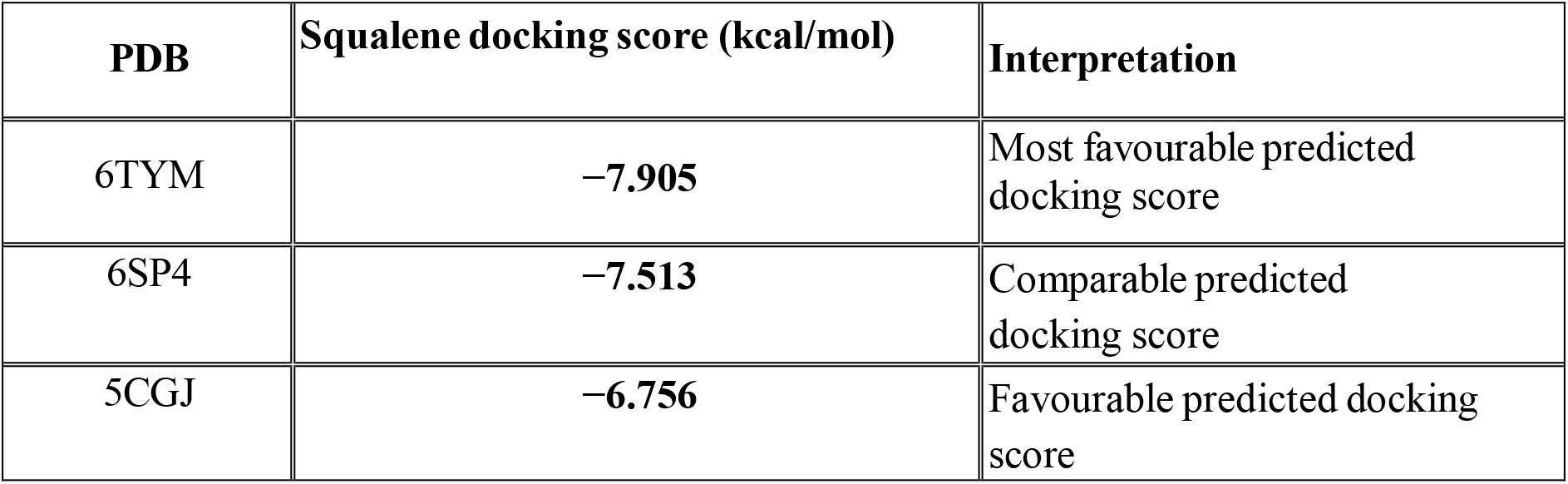
AutoDock Vina docking scores for squalene against validated KEAP1 Kelch-domain crystal structures. Squalene was docked into three independently validated KEAP1 Kelch-domain structures using AutoDock Vina. The reported docking scores correspond to the top-ranked docking poses. The consistency of favourable predicted docking scores across multiple receptor conformations indicates consistent predicted accommodation of squalene within the canonical KEAP1 ligand-binding pocket.

| **PDB** | **Squalene docking score (kcal/mol)** | **Interpretation** |
|---|---|---|
| 6TYM | **−7.905** | Most favourable predicted docking score |
| 6SP4 | **−7.513** | Comparable predicted docking score |
| 5CGJ | **−6.756** | Favourable predicted docking score |

**Figure 6.**
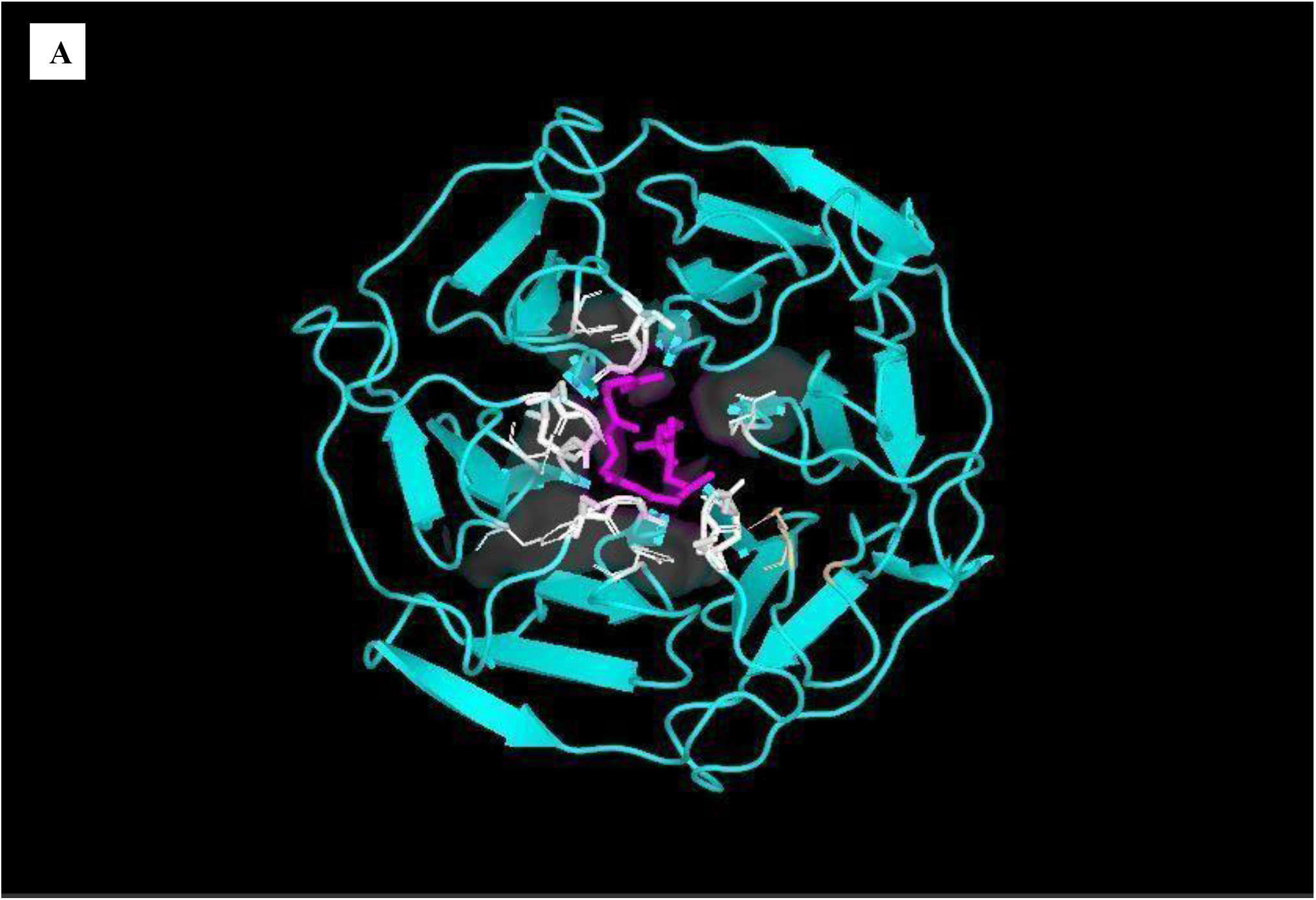
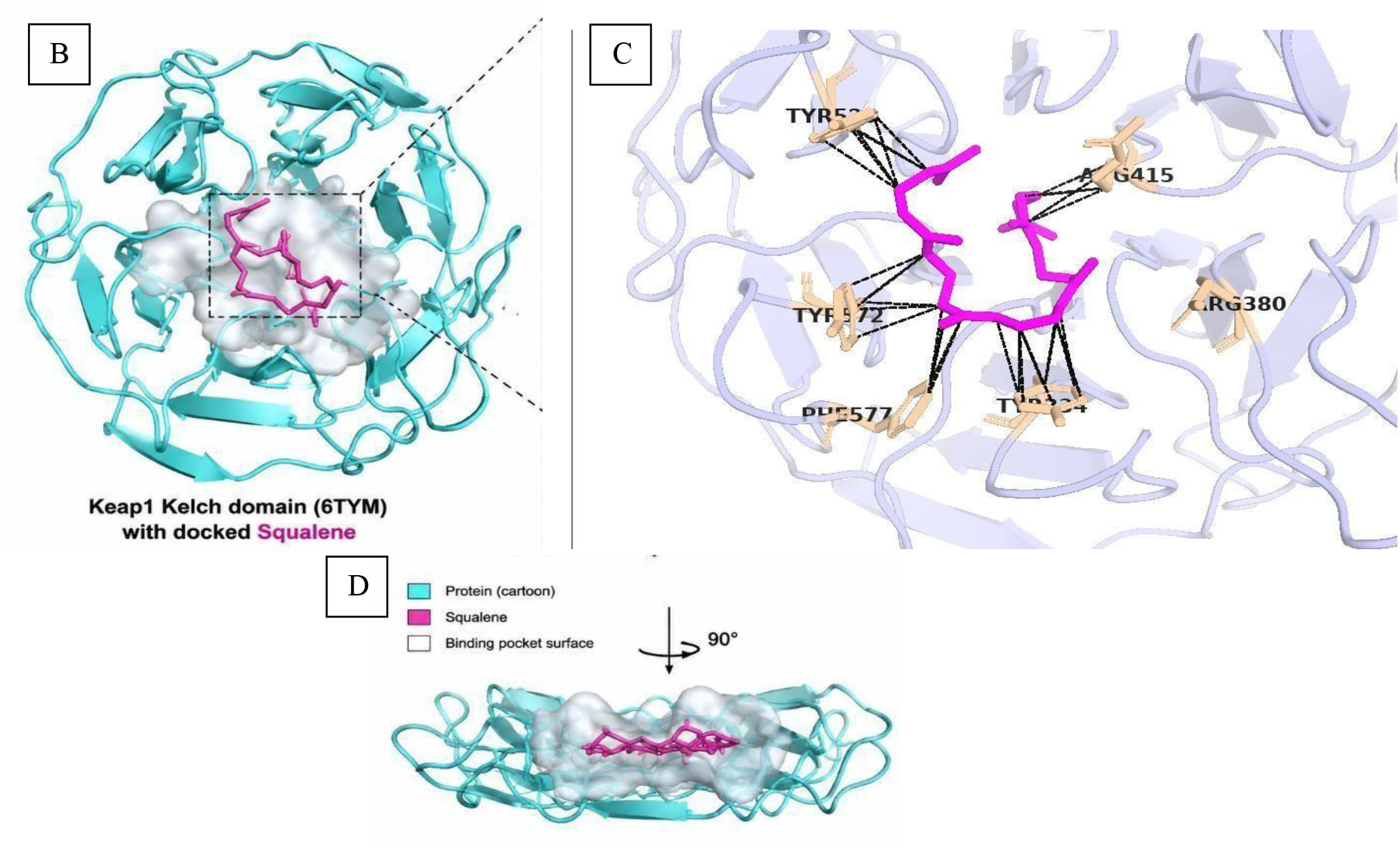
Structural characterisation of the predicted squalene pose within the KEAP1 Kelch domain (PDB ID: 6TYM). Structural representation of the predicted docking pose of squalene within the predefined canonical ligand-binding pocket of the KEAP1 Kelch domain. **(A)** Overall three-dimensional structure showing squalene (magenta) positioned within the predicted binding pocket. **(B)** Surface representation illustrating the accommodation of the predicted squalene pose within the hydrophobic cavity. **(C)** Enlarged view showing the principal residues surrounding the predicted squalene pose. The interaction environment is dominated by hydrophobic and van der Waals contacts in the docking model. **(D)** Orthogonal (90° rotated) view demonstrating the predicted orientation and depth of squalene within the cavity.

**Figure 7.**
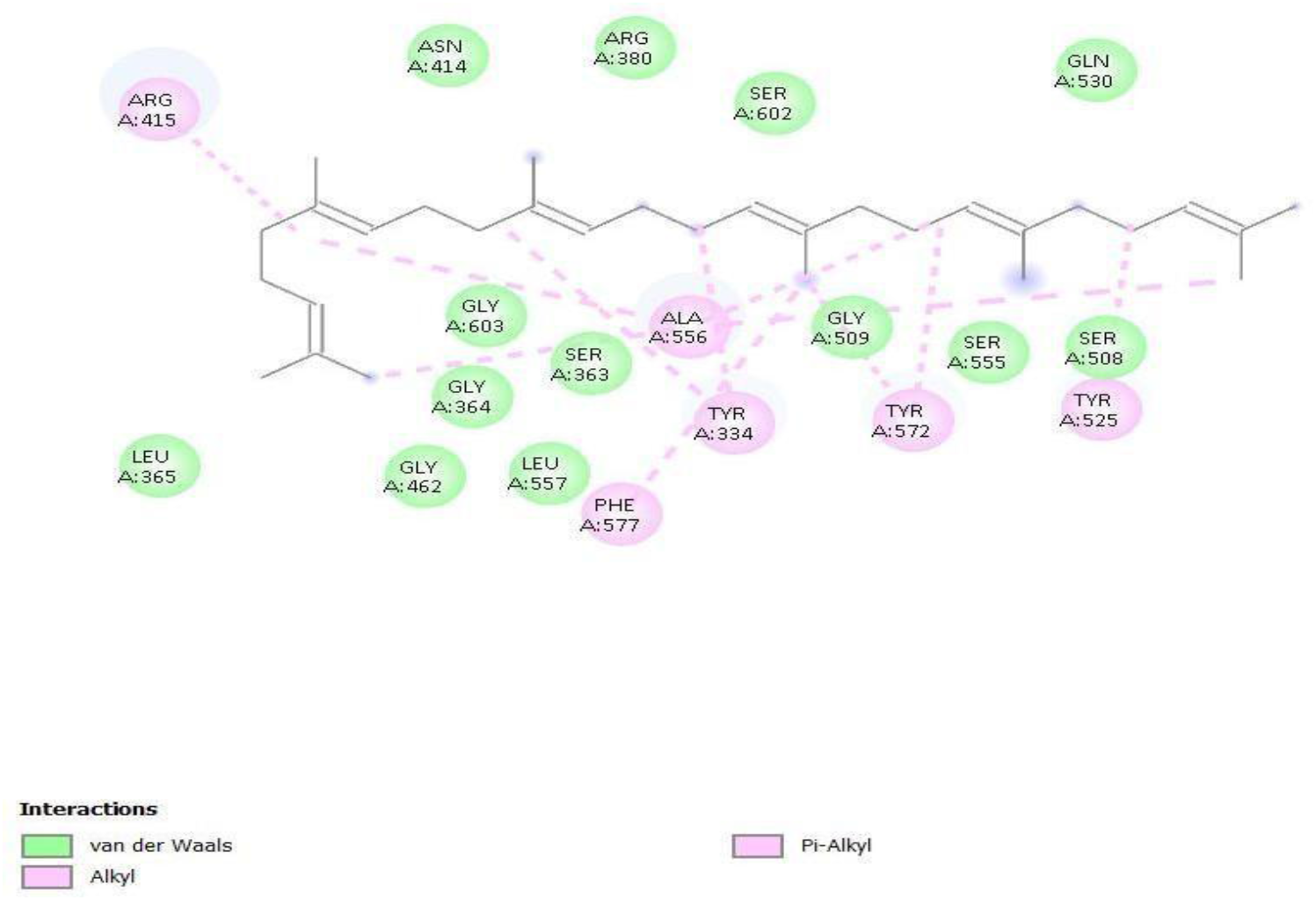
Predicted two-dimensional interaction map of squalene within the KEAP1 Kelch-domain pocket (PDB ID: 6TYM). Two-dimensional representation of the predicted interaction environment generated using Discovery Studio Visualizer. The docking model indicates predominantly hydrophobic and van der Waals contacts involving residues surrounding the predicted squalene pose, including alkyl interactions involving Arg415 and Ala556 and π-alkyl interactions involving Tyr334, Tyr525, Tyr572, and Phe577. Additional surrounding residues contribute through van der Waals contacts. No conventional hydrogen-bond interactions are predicted in the analysed docking pose.

### 3.4 Protein–Ligand Interaction Analysis

Detailed interaction analysis was performed using Discovery Studio Visualizer (BIOVIA, Dassault Systèmes, 2021) to identify the amino acid residues contributing to the stabilisation of the docked squalene molecule within the KEAP1 Kelch domain.

The interaction profile demonstrated that squalene was accommodated within the KEAP1 Kelch-domain binding cavity predominantly through hydrophobic and van der Waals interactions. The principal hydrophobic contacts involved Arg415, Ala556, Tyr334, Tyr525, Tyr572, and Phe577. Specifically, Arg415 and Ala556 contributed alkyl interactions, whereas Tyr334, Tyr525, Tyr572, and Phe577 formed π-alkyl interactions with the squalene molecule. In addition, Arg380, Asn414, Gly509, Ser508, Ser555, Ser602, Gly603, Leu557, and Gln530 were positioned in the surrounding binding environment and contributed predominantly through van der Waals interactions, collectively supporting the positioning and stabilisation of squalene within the binding cavity.

In contrast to the crystallographic ligand, which is capable of forming multiple hydrogen-bond and electrostatic interactions through its polar functional groups, squalene showed no conventional hydrogen-bond interactions in the interaction map and was stabilised primarily through non-polar contacts, including alkyl, π-alkyl, and van der Waals interactions. This interaction pattern is consistent with the highly hydrophobic, linear polyunsaturated structure of squalene, indicating that hydrophobic packing and dispersion-based contacts may play a major role in its accommodation and stabilisation within the KEAP1 Kelch-domain binding pocket (Ferreira et al., 2015; Guedes et al., 2014).

### 3.5 Comparison with the Native Ligand

To compare the predicted interaction environment of squalene with that of the crystallographic ligand, the interaction profile of the docked squalene–KEAP1 complex was compared with that of the crystallographic ligand within the KEAP1 Kelch domain.

The comparison revealed that several residues associated with native ligand recognition, including Arg380, Tyr334, Arg415, Asn414, Tyr572, and Phe577, were conserved within the interaction environment of the docked squalene complex. Although the crystallographic ligand formed hydrogen-bonding and electrostatic interactions that were not observed for squalene, the overlapping interaction environment is consistent with geometrically plausible accommodation of squalene within the predefined canonical KEAP1 pocket. Because the docking grid was centred on the crystallographically defined ligand-binding site, these results do not establish preferential binding to this pocket relative to other regions of the protein.

The distinct interaction pattern observed for squalene can be explained by its chemical composition. Unlike the native ligand, squalene is a non-polar hydrocarbon containing no heteroatoms or polar functional groups capable of forming conventional hydrogen bonds. Consequently, its stabilisation within the binding cavity is expected to depend predominantly on hydrophobic contacts, alkyl and π-alkyl interactions, and van der Waals forces arising from its extensive hydrocarbon surface (Ferreira et al., 2015).

Overall, the conservation of key residues within the binding site, together with the observed hydrophobic interaction profile, provides structural support for the plausible occupation of the canonical KEAP1 Kelch-domain pocket by squalene. However, these docking observations should be considered predictive and do not, by themselves, establish direct binding or functional modulation of KEAP1.

### 3.6 Proposed Mechanistic Model

Based on the computational findings together with previous biological observations from our laboratory, we propose a mechanistic model in which squalene, an evolutionarily conserved isoprenoid, may have a candidate signalling interaction with the KEAP1 Kelch domain. Such an interaction could potentially influence KEAP1–NRF2 signalling and thereby contribute to antioxidant responses. Although this proposed mechanism requires experimental validation, the present computational evidence provides a structural framework for investigating whether the predicted squalene–KEAP1 interaction contributes to previously reported changes in glutathione homeostasis, antioxidant enzyme activity, and stem and progenitor cell protection. If experimentally validated, such an interaction could potentially alter KEAP1–NRF2 recognition and thereby influence NRF2 stabilisation and downstream antioxidant-response signalling (Baird & Yamamoto, 2020; Cuadrado et al., 2019; Kensler et al., 2007).

Although this mechanism requires experimental validation, the present computational evidence provides a structural framework that is consistent with previously reported increases in intracellular glutathione, enhancement of antioxidant enzyme activity, and protection of normal stem and progenitor cells following squalene treatment (Das et al., 2003; Das et al., 2008).

## 4. Discussion

### 4.1 Evolutionary Context and a Testable Co-option Hypothesis

The maintenance of intracellular redox homeostasis represents a fundamental adaptation that enabled aerobic organisms to tolerate the oxidative challenges associated with oxygen metabolism. Antioxidant systems do not simply eliminate reactive oxygen species (ROS), but regulate their abundance within a range compatible with physiological signalling while limiting oxidative damage (Finkel & Holbrook, 2000; Schieber & Chandel, 2014; Sies et al., 2017). The evolutionary conservation of the KEAP1–NRF2 pathway further highlights the importance of regulated antioxidant responses in maintaining cellular stress tolerance (Kensler et al., 2007; Cuadrado et al., 2019; Baird & Yamamoto, 2020).

Squalene biosynthesis is evolutionarily ancient and occurs across diverse prokaryotic and eukaryotic lineages, whereas the canonical KEAP1–NRF2 regulatory system is associated with later-evolving metazoan biology. This temporal distinction does not demonstrate that squalene was historically recruited as a regulator of KEAP1–NRF2 signalling. Nevertheless, it provides a testable evolutionary framework in which an ancient hydrophobic metabolite with membrane-protective and redox-buffering properties could have acquired additional interactions with proteins involved in cellular stress adaptation. The predicted accommodation of squalene within the KEAP1 Kelch-domain cavity is compatible with this possibility, but molecular docking cannot establish evolutionary co-option, physiological binding, or functional pathway regulation.

In the present study, the initial MOE screening identified KEAP1 as the most favourable target among the selected proteins evaluated under the applied workflow. This analysis was used for target prioritisation rather than as a quantitative comparison of binding affinities between unrelated proteins and binding sites. Subsequent validation using three independent KEAP1 structures—6TYM, 6SP4, and 5CGJ—demonstrated RMSD values below 2.0 Å, supporting the ability of the docking protocol to reproduce experimentally determined ligand conformations.

Importantly, native-ligand redocking validates the computational protocol’s ability to reproduce known ligand-binding geometries; it does not establish that squalene binds KEAP1 under physiological conditions. The present results should therefore be interpreted as a structural hypothesis supporting further investigation of a possible squalene–KEAP1 interaction rather than definitive evidence of KEAP1-mediated signalling. Linking Computational Predictions with previous biological evidence

One of the major strengths of the present work is that previous biological observations from our laboratory support the computational findings. Earlier investigations demonstrated that squalene selectively protects normal hematopoietic and mesenchymal stem and progenitor cells against platinum-induced oxidative injury while preserving intracellular glutathione homeostasis and enhancing glutathione-dependent antioxidant enzyme activity (Das et al., 2003; Das et al., 2008). These biological effects could not be fully explained by the direct free-radical scavenging properties of squalene alone, suggesting the involvement of upstream regulatory signalling pathways (Popa et al., 2015; Lozano-Grande et al., 2018).

The present docking results provide a plausible molecular framework that may explain these earlier observations. The predicted interaction of squalene with the KEAP1 Kelch domain raises the possibility that squalene may indirectly promote NRF2 activation, leading to increased expression of genes involved in glutathione biosynthesis, antioxidant defence, and cellular stress adaptation (Kensler et al., 2007; Cuadrado et al., 2019; Baird & Yamamoto, 2020). Very recently, independent experimental findings have strengthened the biological basis for linking squalene to NRF2-dependent antioxidant regulation. Jiang et al. (2026) demonstrated that squalene increased Nrf2 and downstream HO-1 and NQO1 expression in DSS-induced colitis, while genetic Nrf2 deficiency markedly attenuated the antioxidant, anti-inflammatory and tissue-protective effects of squalene. Pharmacological inhibition of Nrf2 similarly reduced squalene-mediated protection in Caco-2 cells. Furthermore, CETSA indicated increased thermal stability of Nrf2 following squalene treatment, while molecular docking and 100-ns molecular-dynamics simulations supported a potential direct squalene–Nrf2 interaction. These findings provide independent evidence that the biological effects of squalene can be NRF2-dependent. However, they do not establish whether direct Nrf2 interaction is the sole mechanism by which squalene engages this pathway. The present prediction of squalene accommodation within the KEAP1 Kelch domain therefore raises a complementary hypothesis in which squalene may influence the NRF2 regulatory axis at more than one molecular level. Definitive characterization of direct squalene–Nrf2 binding and binding kinetics will require orthogonal biophysical approaches. Although direct experimental evidence remains to be established, this hypothesis is consistent with the previously reported enhancement of glutathione metabolism observed following squalene treatment (Das et al., 2008).

The present study proposes a distinct mechanism whereby oxidative stress may recruit a KEAP1–NRF2-dependent antioxidant program through the interaction of squalene with the KEAP1 Kelch domain. Together, these observations suggest that different stress-responsive pathways may contribute to stem-cell adaptation (stem cell altruism), although whether KEAP1–NRF2 signalling participates in this process following squalene exposure remains to be experimentally determined. Different environmental challenges may recruit distinct molecular pathways that ultimately converge on the same adaptive biological outcome—temporary enhancement of stem cell robustness, preservation of the surrounding niche, and maintenance of tissue homeostasis.

### 4.2 Squalene as a Stress-responsive Isoprenoid with Potential Signalling Function

Traditionally, squalene has been regarded as a biosynthetic intermediate of the mevalonate pathway whose primary biological function is to serve as a precursor for sterol synthesis (Spanova & Daum, 2011; Goldstein & Brown, 1990). However, this interpretation may underestimate its broader biological significance. Squalene is evolutionarily conserved across diverse kingdoms of life, possesses a unique linear thirty-carbon hydrocarbon backbone, and exhibits physicochemical properties that distinguish it from downstream sterols such as lanosterol and cholesterol (Spanova & Daum, 2011; Popa et al., 2015; Hazra & Chattopadhyay, 2023).

From an evolutionary perspective, terpenoid metabolites have been proposed to represent ancient components of cellular biochemistry, providing a broader context for considering whether conserved isoprenoids may possess functions beyond their canonical metabolic roles (Ourisson & Nakatani, 1994). We propose that squalene may have biological functions beyond its established role as a metabolic intermediate of the mevalonate pathway. Its evolutionary conservation and highly flexible hydrophobic structure provide a rationale for investigating whether it can participate in protein–lipid interactions with signalling proteins (Wang & Casey, 2016; Spanova & Daum, 2011). Recent evidence further supports the concept that squalene may function as a stress-responsive metabolite rather than solely as a passive biosynthetic intermediate. Squalene monooxygenase (SQLE) catalyses the conversion of squalene to 2,3-oxidosqualene, and hypoxia-associated alterations in SQLE activity can promote intracellular squalene accumulation (Coates et al., 2023), whereas reduced SQLE activity and consequent squalene accumulation have independently been shown to alter membrane lipid composition, reduce lipid peroxidation, and decrease susceptibility to ferroptotic cell death (Garcia-Bermudez et al., 2019). Taken together, these observations raise the testable hypothesis that hypoxia-associated squalene accumulation may contribute to an endogenous membrane-protective response during oxygen limitation. Importantly, such a membrane-associated protective function would be mechanistically distinct from, but potentially complementary to, the predicted interaction of squalene with the KEAP1 Kelch domain investigated in the present study.

Viewed from a systems-biology perspective, hypoxia-associated squalene accumulation may represent a molecular example consistent with the previously proposed concept of adaptive metabolic disequilibrium. In this framework, environmental stress transiently displaces metabolic flux from its preceding steady state, generating a new metabolic configuration that may itself confer adaptive value. Under hypoxia, altered oxygen-dependent SQLE activity may increase intracellular squalene availability, and the accumulated metabolite may contribute to membrane protection against lipid peroxidation and ferroptotic injury. Thus, the stress-induced alteration of sterol-pathway flux may not represent only metabolic blockade, but potentially an adaptive redistribution of metabolic resources that enhances cellular robustness. Whether this phenomenon constitutes a general mechanism of adaptive disequilibrium remains to be established experimentally.

An important consideration for this proposed signalling mechanism is the membrane association and protein accessibility of squalene. Given its highly hydrophobic structure, squalene is expected to partition preferentially into lipid-rich environments, raising the question of whether a sufficient fraction becomes accessible for interaction with soluble proteins such as the KEAP1 Kelch domain. The present docking analysis does not address the subcellular distribution, membrane partitioning, or effective concentration of protein-accessible squalene and therefore cannot establish whether the predicted KEAP1 interaction occurs under physiological conditions.

Nevertheless, several potential routes could permit transient encounters between membrane-associated squalene and cytosolic proteins. Squalene may become transiently accessible at ER–cytosol interfaces or at the surface of lipid droplets, where active lipid exchange and membrane remodelling could facilitate movement between membrane-associated and protein-accessible pools. Alternatively, intracellular lipid-transfer or sterol-handling proteins could potentially facilitate the redistribution or presentation of hydrophobic metabolites between lipid compartments and soluble protein environments. Such mechanisms would not require squalene to exist as a freely soluble metabolite in the aqueous cytosol; rather, transient local availability at membrane interfaces or within protein-associated lipid-transfer pathways could provide opportunities for interaction with cytosolic signalling proteins.

This model is particularly relevant to the present docking prediction because the KEAP1 Kelch-domain cavity itself is predominantly hydrophobic, and the predicted squalene interaction is mediated largely through hydrophobic and van der Waals contacts rather than conventional hydrogen bonding. Thus, the physicochemical properties that favour squalene partitioning into lipid environments may also contribute to its compatibility with a hydrophobic protein-binding cavity. However, structural compatibility should not be equated with physiological accessibility. Direct experimental determination of squalene localisation, membrane partitioning, protein accessibility, and potential lipid-transfer mechanisms will be required to establish whether sufficient squalene reaches KEAP1 in living cells.

The present docking results therefore provide a structural hypothesis for a possible squalene–KEAP1 interaction rather than evidence of physiological binding. If experimentally validated, transient mobilisation of squalene from membrane-associated pools could provide a mechanistic link between metabolic stress adaptation, membrane protection, and protein-mediated antioxidant signalling. Such a mechanism could expand current understanding of the biological functions of the mevalonate pathway while providing a testable connection between squalene metabolism and KEAP1–NRF2-dependent cellular stress responses.

### 4.3 Rotational Flexibility and Interpretation of the Squalene Docking Result

An important consideration in interpreting the present docking results is the unusually high conformational flexibility of squalene. Unlike the relatively constrained small-molecule ligands used for native-ligand redocking, squalene possesses a long, acyclic polyisoprenoid backbone with numerous potentially rotatable single bonds. Consequently, its conformational search space is substantially larger than that of the crystallographic ligands used for protocol validation.

This distinction introduces an important limitation when comparing the confidence of squalene docking with the excellent native-ligand RMSD values obtained during validation. The RMSD values of 0.826 Å, 0.277 Å, and 0.303 Å for 6TYM, 6SP4, and 5CGJ, respectively, demonstrate that the selected docking protocol can reproduce the experimentally observed conformations of the corresponding crystallographic ligands. However, successful redocking of relatively constrained ligands does not necessarily demonstrate equivalent conformational sampling of a highly flexible molecule such as squalene. Accordingly, the predicted squalene docking scores of −7.905, −7.513, and −6.756 kcal/mol for 6TYM, 6SP4, and 5CGJ, respectively, should be regarded as comparative docking scores rather than experimental estimates of binding free energy. The elongated hydrocarbon chain can adopt multiple conformations within a hydrophobic cavity, and different conformers may produce similar docking scores while differing substantially in their detailed orientations.

The final squalene PDBQT contained 15 torsional degrees of freedom (TORSDOF = 15). For subsequent squalene docking, AutoDock Vina was run with an exhaustiveness of 64 to provide broader conformational sampling of the highly flexible ligand.

This limitation does not invalidate the observed structural prediction, but it reinforces the need for additional conformational sampling, replicate docking, molecular dynamics simulations, and ultimately experimental binding measurements. The present results therefore identify a plausible binding mode rather than establishing a unique or energetically definitive squalene conformation.

### 4.4 Broader Implications for Stem-Cell Niche Defence and Adaptive Altruism

Stem-cell survival depends on the maintenance of intracellular redox homeostasis, in which controlled reactive oxygen species (ROS) signalling supports self-renewal and regenerative potential, whereas excessive oxidative stress can contribute to stem-cell exhaustion (Bigarella et al., 2014; Ito & Suda, 2014; Schieber & Chandel, 2014). Previous studies from our laboratory demonstrated that squalene enhances intracellular glutathione levels and protects normal stem and progenitor cells against oxidative injury (Das et al., 2003; Das et al., 2008). More recent experimental work has shown that mesenchymal stem cells can exert measurable cytoprotective effects against platinum-mediated injury, including preservation of colony formation in cisplatin-exposed hematopoietic stem/progenitor cells, providing an experimental basis for considering cytoprotection as a component of stem-cell niche defence (Sandhya et al., 2026). Together, these findings raise the possibility that squalene-mediated redox protection may contribute to cellular stress adaptation and, potentially, to broader stem-cell niche defence under adverse environmental conditions.

This possibility can be considered within the broader Adaptive Altruism framework, which proposes that transient stress-adapted stem-cell states may acquire enhanced metabolic fitness and stress resistance before potentially contributing to the preservation of the surrounding cellular niche (Talukdar et al., 2026). Recent findings further suggest that squalene may participate in stress adaptation through a membrane-associated anti-ferroptotic mechanism. Hypoxia can promote intracellular squalene accumulation through oxygen-sensitive regulation of SQLE (Coates et al., 2023), while reduced SQLE activity and consequent squalene accumulation have been associated with altered membrane lipid composition, reduced lipid peroxidation, and decreased susceptibility to ferroptotic oxidative injury (Garcia-Bermudez et al., 2019). Such membrane-associated protection may represent an immediate physicochemical component of cellular stress resistance. In parallel, the present docking findings raise a distinct but potentially complementary hypothesis in which a protein-accessible fraction of squalene interacts with the KEAP1 Kelch domain and thereby influences NRF2-associated antioxidant signalling. If such an interaction results in NRF2 activation, increased expression of antioxidant and glutathione-associated genes, including GCLC, GCLM and SLC7A11, could enhance cellular glutathione availability and support GPX4-dependent detoxification of lipid hydroperoxides. Thus, squalene-mediated membrane protection and KEAP1–NRF2-associated antioxidant signalling may represent mechanistically distinct but potentially coordinated components of cellular defence against oxidative and lipid-peroxidative stress.

The concept of altruistic stem cells (ASCs) proposes that transient stress-adapted stem-cell states may contribute to protection of the surrounding niche through enhanced cellular robustness and cytoprotective mechanisms (Das et al., 2012; Pathak et al., 2021). This concept has subsequently contributed to the broader Adaptive Altruism framework, in which transient stress-adapted cellular states are proposed to acquire enhanced metabolic fitness and stress resistance before potentially contributing to collective niche protection (Talukdar et al., 2026). The earlier concept of organised adaptive disequilibrium provides a systems-level context for this transition, in which environmental stress may transiently reorganise metabolic and signalling networks to generate a state with increased cellular robustness rather than merely restoring the preceding steady state. The historical and conceptual development of this framework has been described in The Science Behind Squalene (Das, 2000) and its accompanying commentary.

Within this broader Adaptive Altruism framework, the hypoxia–SQLE–squalene response described by Coates et al. (2023) may provide a candidate metabolic example of the proposed initial enhanced-fitness phase of stress-adapted stem or progenitor cells. Transient metabolic disequilibrium could increase cellular robustness through squalene accumulation, preservation of membrane integrity, reduction of lipid peroxidation, and maintenance of antioxidant capacity. If the predicted KEAP1–NRF2 interaction is subsequently validated, NRF2-dependent enhancement of glutathione metabolism could provide an additional layer of protection against oxidative stress. Together, these effects could generate a transiently stress-resistant cellular state that is better equipped to tolerate an adverse microenvironment.

A further possibility is that such stress-adapted cells could subsequently contribute to niche protection. Within the proposed altruistic stem-cell “helper” phenotype, enhanced fitness of transient actor cells precedes collective protection of neighbouring cells. In this model, increased antioxidant capacity in the actor-cell population could theoretically reduce the local burden of reactive or lipid-oxidising species and thereby contribute to preservation of neighbouring stem or progenitor cells. This possibility is particularly relevant to environments characterised by substantial oxidative stress, including exposure to platinum-based chemotherapeutic agents, where selective protection of normal stem and progenitor populations has previously been observed following squalene treatment (Das et al., 2003; Das et al., 2008).

Our computational findings provide a structural mechanism for what was originally conceptualized as an adaptive mechanism under metabolic stress and chaos, governed by the force of disequilibrium (Das, 2000). In the context of the Tumor Stemness Switch (TSS), transient hypoxia downregulates SQLE, thereby shifting the cellular operating point. The resulting squalene accumulation may contribute to membrane protection against lipid peroxidation and, if a KEAP1 interaction is experimentally validated, could provide an additional upstream regulatory influence on glutathione synthesis and export relevant to non-autonomous niche protection.

Accordingly, the present findings suggest a testable mechanistic continuum linking metabolic stress adaptation with the proposed Adaptive Altruism framework: **environmental stress → altered SQLE-dependent metabolic flux → squalene accumulation → membrane protection and reduced lipid peroxidation → potential KEAP1–NRF2 activation → enhanced glutathione-dependent antioxidant capacity → increased stress-adapted cellular fitness → potential niche-supportive/helper-cell phenotype**. This model integrates the membrane-protective properties of squalene with its proposed protein-mediated signalling function and provides a conceptual framework connecting metabolic adaptation, redox regulation, and stem-cell niche defence. However, each step downstream of squalene accumulation remains to be experimentally established. The present study does not demonstrate that squalene activates NRF2, increases GCLC, GCLM or SLC7A11 expression, enhances GPX4-dependent lipid peroxide detoxification through KEAP1, or induces a helper-cell phenotype. Likewise, it does not establish that squalene-treated cells alter the oxidative microenvironment sufficiently to protect neighbouring stem cells, or that such protection preserves stem-cell quiescence or genomic integrity. These relationships therefore remain testable hypotheses requiring direct biochemical, cellular, and co-culture studies.

### 4.5 Limitations and Future Perspectives

Several limitations should be considered when interpreting the present findings. First, molecular docking provides a static representation of protein–ligand recognition and cannot establish physiological binding or signalling activity. Although native-ligand redocking produced excellent RMSD values across three KEAP1 structures, these results primarily validate the docking protocol and do not demonstrate that squalene binds KEAP1 in living cells.

Second, the highly flexible nature of squalene introduces greater conformational uncertainty than that associated with the crystallographic ligands used for validation. Explicit reporting of the number of active rotatable bonds and the squalene-specific docking exhaustiveness is therefore important for reproducibility. Additional replicate docking, expanded conformational sampling and molecular dynamics simulations will be required to determine whether the predicted binding orientation remains stable over time.

Third, the present computational workflow does not address the intracellular accessibility of squalene. Its preferential association with membranes, lipid droplets and other hydrophobic compartments raises the possibility that only a fraction of the cellular squalene pool may be available for interaction with cytosolic KEAP1. Future studies should therefore investigate intracellular localisation, lipid partitioning and possible lipid-transfer mechanisms in parallel with direct binding measurements.

Because squalene is highly hydrophobic, uniform aqueous cytosolic distribution is physically improbable. However, consistent with the compartmentalized squalene pools described in our foundational work (Das, 2000), squalene accumulation occurs prominently at endoplasmic reticulum (ER) and lipid droplet membranes. Given that KEAP1 scans substrate pools in close proximity to the ER-cytosol interface, localized interphase partitioning could satisfy the concentration threshold required for squalene to engage the Kelch propeller without requiring bulk aqueous solubility.

Finally, the proposed connection between squalene, KEAP1–NRF2 signalling, ferroptosis resistance and Adaptive Altruism requires experimental validation at multiple biological levels. Biophysical experiments should determine whether squalene directly associates with the KEAP1 Kelch domain. Cellular studies should examine NRF2 stabilisation, nuclear localisation, ARE-dependent transcription, GCLC/GCLM/SLC7A11 expression, glutathione levels and GPX4-dependent lipid peroxide detoxification following squalene exposure. Ferroptosis assays should determine whether these responses are functionally linked to resistance against lipid peroxidation. In stem and progenitor cell systems, co-culture and conditioned-medium experiments could further test whether stress-adapted cells acquire a measurable niche-supportive phenotype.

A further priority is rigorous molecular dynamics analysis of the predicted squalene–KEAP1 complex. Future simulations should include independent replicate trajectories to evaluate complex stability, ligand retention, residue-level contact occupancy, protein flexibility, and solvent exposure under physiologically relevant conditions. Binding free-energy estimation and comparison with native KEAP1 ligands could further assess the relative stability and plausibility of the predicted interaction. These computational analyses should be complemented by biophysical binding assays and cellular studies of NRF2 activation, glutathione metabolism, lipid peroxidation, and ferroptosis-related responses.

If these predictions are experimentally validated, the resulting model would position squalene not simply as a passive intermediate of sterol biosynthesis, but as a metabolically responsive isoprenoid capable of integrating membrane protection with antioxidant signalling and potentially contributing to multicellular stress adaptation.

## 5 Conclusion

This study builds on earlier work on squalene metabolism and its possible role in protecting cells under stress (Das, 2000; Das et al., 2012; Pathak et al., 2025; Talukdar et al., 2026). Our computational analysis suggests that squalene can fit within the ligand-binding pocket of the KEAP1 Kelch domain. The predicted interaction is mainly supported by hydrophobic and van der Waals contacts with residues in the binding pocket. Native-ligand redocking also reproduced the experimentally observed ligand positions with RMSD values below 1 Å, supporting the reliability of the docking procedure.

These findings raise the possibility that squalene may have a role beyond its established function in cellular metabolism. Previous studies have reported protective effects of squalene, including its association with glutathione homeostasis, antioxidant responses, and stem-cell protection (Das et al., 2008; Baird & Yamamoto, 2020; Cuadrado et al., 2019). The predicted interaction with KEAP1 provides a possible link between these observations and the KEAP1–NRF2 pathway, which regulates cellular antioxidant responses. However, whether squalene directly interacts with KEAP1 and influences this pathway remains to be established.

Further studies, including molecular dynamics simulations, biophysical binding experiments, intracellular localisation studies, and cellular assays, are needed to investigate this possibility. These studies should determine whether squalene interacts with KEAP1 under physiological conditions and whether this interaction contributes to NRF2 activation and cellular protection. Overall, the present work provides a computational structural hypothesis for a possible squalene–KEAP1 interaction; it does not establish direct physiological binding or functional NRF2 activation.

## 6. Data Availability

All computational data generated and analysed during this study, including Molecular Operating Environment (MOE) target-screening files, receptor and ligand structures, receptor and ligand PDBQT files, AutoDock Vina configuration files, docking output pose files, native-ligand redocking validation files, and protein–ligand interaction analysis files, are publicly available in the Zenodo repository: https://doi.org/10.5281/zenodo.22942431 The MOE database files require compatible MOE software for direct inspection.

## 7. Author Contributions

PD performed the molecular docking and computational analyses, conducted native-ligand redocking validation using AutoDock Vina, analysed protein–ligand interactions, curated the data, prepared the figures and tables, and drafted the manuscript. BD conceptualised and supervised the study, contributed to the interpretation of the computational findings, proposed the biological mechanisms, and critically reviewed and edited the manuscript. LP,RD, PJS, DD, SB, and US critically reviewed the manuscript, provided editorial input, and contributed relevant literature and references to support the scientific discussion. All authors reviewed and approved the final version of the manuscript.

## 8. Acknowledgements

We sincerely extend our gratitude to the members of KaviKrishna Laboratory, Research Park, Indian Institute of Technology Guwahati, Guwahati, Assam, India, and the Thoreau Laboratory for Global Health, M2D2, University of Massachusetts, Lowell, MA, USA, for their valuable insights, support, and expertise, which greatly contributed to the completion of this research article. We also express our sincere gratitude to the members of KaviKrishna Telemedicine Care, Sualkuchi, Assam, India, for their continued support and encouragement.

## 10. Competing Interests

The authors declare no competing financial or non-financial interests.

Expression of CD271, CD133, CD45, CD105, CD73, CD90, VEGFR2, ABCG2 and HIF-2α, together with CFU-F frequency, is compared between serum-free and serum-containing culture conditions. Oct-4 expression was assessed by qPCR. Values are presented as reported in the original source (mean ± SD; n = 5 for phenotypic analysis). CFU-F values are expressed per 10⁶ assayed cells.

---

**Supplementary Table S1.** AutoDock Vina docking parameters used for native-ligand validation and squalene docking.

| **Receptor** | **Ligand** | **X** | **Y** | **Z** | **Size X** | **Size Y** | **Size Z** | **Exhaust.** | **Num. modes** | **Energy range** | **Seed** |
|---|---|---|---|---|---|---|---|---|---|---|---|
| **6TYM** | Native ligand 08A | −5.021 | −23.723 | −19.404 | 22.5 | 22.5 | 22.5 | **8** | 10 | 3 (default) | Not specified |
| **6SP4** | Native ligand LQK | 36.511 | 3.243 | 24.143 | 22.5 | 22.5 | 22.5 | **32** | 20 | 3 (default) | Not specified |
| **5CGJ** | Native ligand 51M | 39.832 | −20.257 | −4.736 | 22.5 | 22.5 | 22.5 | **32** | 20 | 3 (default) | Not specified |
| **6TYM** | Squalene | −5.021 | −23.723 | −19.404 | 22.5 | 22.5 | 22.5 | **64** | 20 | **5** | Not specified |
| **6SP4** | Squalene | 36.511 | 3.243 | 24.143 | 22.5 | 22.5 | 22.5 | 64 | 20 | 3 (default) | Not specified |
| **5CGJ** | Squalene | 39.832 | −20.257 | −4.736 | 22.5 | 22.5 | 22.5 | 64 | 20 | 3 (default) | Not specified |

Table Note: Coordinates are in Å. “3 (default)” indicates that the Vina default energy range was used because no explicit value was specified in the configuration file. Random seeds were not explicitly specified in the recorded configurations.

**Supplementary Table S2.** Protein structures used for preliminary MOE target screening.

| Target protein | PDB ID | Structural region/domain | Docking site | Reason for inclusion |
|---|---|---|---|---|
| **KEAP1** | 1U6D | Kelch domain | Ligand-binding pocket | Antioxidant/NRF2 signalling |
| **HMG-CoA reductase** | 1DQ8 | Catalytic domain | Ligand-binding region | Mevalonate/cholesterol biosynthesis |
| **c-Myc** | 1NKP | Myc–Max DNA-binding complex | Selected protein surface/site used in MOE | Cell proliferation/stress-related signalling |
| **p53** | 1TUP | DNA-binding domain | Selected site used in MOE | Stress-response pathway |

**Table S1.** Phenotypic characterization and colony-forming capacity of bone marrow mesenchymal stromal cells maintained under serum-free hypoxic suspension culture conditions.

| **MSC phenotype assessment** | **CD271+/CD133+**<br>**serum free** | **CD271+/CD133+**<br>**20% FBS** |
|---|---|---|
| CD271 | 92.7 ± 3.6 | 11 ± 4.3 |
| CD133 | 73 ± 8.5 | 8 ±2.1 |
| CD45 | 24 ± 9.7 | 9.3 ± 2.1 |
| CD105 | 47.7 ± 12.2 | 84.4 ± 21.8 |
| CD73 | 56.3 ± 8.7 | 75 ± 14.4 |
| CD90 | 21 ± 6.5 | 73.5 ± 15.2 |
| VEGFR2 | 2 ± 1.2 | 1.2 ± 0.4 |
| ABCG2 | 24 ± 19.6 | 1.1 ± 0.4 |
| HIF-2α | 20± 17.2 | 2.1 ± 1.4 |
| Oct-4 | - | - |
| CFU-F/10<sup>6</sup> cells) | 4349 ± 1936 | 3910 ±1993 |

(\*) CD271+/CD133+ cells were cultured for two weeks with or without serum plus TPO, Flt3 ligand and SCF and subjected to flow cytometry.  Note that about 92% of cells remained positive for CD271 in the serum-free culture system whereas only 11% cells remain CD271+ when grown in the presence of serum (details given in method).  Oct-4 values were obtained by QPCR.  n=5

Expression of CD271, CD133, CD45, CD105, CD73, CD90, VEGFR2, ABCG2 and HIF-2α, together with CFU-F frequency, is compared between serum-free and serum-containing culture conditions. Oct-4 expression was assessed by qPCR. Values are presented as reported in the original source (mean ± SD; n = 5 for phenotypic analysis). CFU-F values are expressed per 10^6^ assayed cells.

**Data provenance:** These data were previously published in [insert the verified original reference]. They are reproduced or adapted here with appropriate attribution and any required permission or licence.

